# Device-embedded accelerometry complements neural signals for tracking parkinsonian motor states

**DOI:** 10.64898/2026.07.08.737286

**Authors:** Tao Liu, Jiaang Yao, Bahman Abdi-Sargezeh, Abhinav Sharma, Camille Lasbareilles, Robert Tsi Lok Ho, Jackson T.S. Cheung, Timothy Denison, Huiling Tan, Wolf-Julian Neumann, Minghan Max Zhu, Sebastian Liu, Philip A. Starr, Simon Little, Ashwini Oswal

## Abstract

Adaptive deep brain stimulation (aDBS) relies on physiological biomarkers to infer motor state and guide therapeutic stimulation in Parkinson’s disease. However, neural biomarkers may themselves be altered by stimulation, potentially limiting their utility for closed-loop control. We address this limitation by testing whether DBS device-embedded accelerometers can accurately track Parkinsonian motor state across stimulation conditions. We analysed over 1,900 hours of chronic recordings of subthalamic nucleus (STN), sensorimotor cortical and device-embedded accelerometry signals acquired before and during continuous STN stimulation, alongside continuous wearable assessments of bradykinesia and dyskinesia. Across stimulation conditions, accelerometry-derived features robustly tracked motor symptom severity and outperformed neural features for symptom decoding. Mechanistically, total STN beta power - a widely used biomarker for aDBS - proved less informative because it conflates periodic and aperiodic neural processes with opposing relationships to motor state. Under active stimulation, periodic beta activity showed reduced coupling to symptom severity, whereas STN aperiodic activity, cortical periodic activity and cortico-subthalamic coherence remained comparatively stable. Together, these findings demonstrate that neural and behavioural biomarkers exhibit differential robustness during deep brain stimulation and identify device-embedded accelerometry as a robust behavioural biomarker of motor state, motivating its use in next-generation adaptive DBS systems.

## Introduction

Bradykinesia is a defining motor symptom of Parkinson’s disease (PD), characterized by abnormally reduced movement speed and amplitude, which severely impairs daily function and quality of life^1^. Although dopaminergic therapy effectively mitigates bradykinesia, chronic treatment often induces involuntary hyperkinetic movements known as dyskinesias^2–4^.

Deep Brain Stimulation (DBS) of the subthalamic nucleus (STN) is an effective treatment for medication refractory bradykinesia that also ameliorates dyskinesia by facilitating reductions in dopaminergic medication dosage^5–8^. Despite its success, DBS has yet to reach its full potential for personalised therapy. Conventional DBS systems deliver continuous, high frequency stimulation pulses that ignore moment-to-moment fluctuations in neural dynamics which govern clinical state. Adaptive (closed loop) DBS offers refinement by titrating stimulation to clinical states inferred from physiological biomarkers - delivering more stimulation when pathological (bradykinetic) activity is detected and less when it is not. Importantly, because adaptive DBS relies on biomarkers measured during active stimulation, candidate control signals must remain informative despite the very intervention they are intended to guide. In this regard, beta band (13-30 Hz) activity within the STN has been the most extensively studied biomarker of bradykinesia and rigidity^9–14^ - assessed using part III of the Movement Disorder Society Unified Parkinson’s Disease Rating Scale (UPDRS)^15^ - leading to its widespread use as a control signal for adaptive DBS^16,17^. Yet, total beta power conflates two distinct physiological processes: a periodic (oscillatory) component reflecting rhythmic firing^18,19^, and an aperiodic component linked to excitation-inhibition balance in STN spiking activity^20–23^. How these components, and other spectral activities within the cortico-STN motor network, dynamically relate to bradykinesia and dyskinesia remains unresolved, largely because standard clinical assessments cannot capture motor state fluctuations at the timescale of neural dynamics. This highlights an urgent need for objective, high temporal resolution biomarkers capable of continuously quantifying motor state^24–26^.

Wrist-worn accelerometers have shown strong potential for estimating clinical states^2,27–29^, inspiring commercial monitoring systems such as the Parkinson’s KinetiGraph (PKG)^29,30^. More recently, sensors placed at alternative body sites, including the ankle and waist, have shown comparable efficacy for tracking symptom dynamics^31,32^. The use of wearable technology biomarkers for adaptive DBS poses a challenge however, due to both the potential for user non-compliance and the need for real-time integration of signals between the wearable and the implantable pulse generator (IPG, which is often located within the chest wall). A promising alternative lies in the accelerometers already embedded in many modern DBS IPGs^33–35^, which could provide continuous, implant-based behavioral readouts without external hardware. However, it remains unknown whether IPG accelerometry can reliably capture fluctuations in bradykinesia and dyskinesia severity with sufficient sensitivity and specificity for adaptive control.

To address these questions, we recorded over 1900 hours of simultaneous sensorimotor cortical, STN, and device accelerometer data from individuals with Parkinson’s disease implanted with an investigational sensing-enabled DBS system (986 hours without stimulation, 915 hours with continuous stimulation). Concurrent measurements of bradykinesia and dyskinesia severity were continuously quantified using bilaterally worn PKG devices, whilst participants performed activities of daily living prior to and after the initiation of therapeutic DBS. These multimodal recordings enabled us to: **(i)** identify neural features within the motor cortical–STN circuit that best predict bradykinesia and dyskinesia severity, **(ii)** evaluate the capacity of device-embedded accelerometry to capture these motor fluctuations, and **(iii)** determine whether integrating neural and accelerometry features improves clinical state estimation relative to either modality alone.

## Materials and Methods

### Patient characteristics and surgical procedure

We studied 11 patients diagnosed with idiopathic Parkinson’s disease, who underwent bilateral implants of the investigational Summit RC + S neural interface (Medtronic). Participants were recruited from surgical movement disorders clinics at the University of California San Francisco and had standard clinical indications for STN DBS^36^. All patients provided written, informed consent and study procedures were approved by the institutional review board. Clinical characteristics of the patients are provided in **Table 1**. A movement disorders specialist assessed baseline motor function using the Movement Disorder Society Unified Parkinson’s Disease Rating Scale part III (UPDRS III) in both OFF- and ON-medication states before implantation.

**Table 1:** Patient demographics and clinical characteristics.

| Patient | Age<br>(years) | Gender | Disease<br>duration<br>(years) | Preoperative<br>medication<br>(mg) | UPDRS<br>III<br>(off/on) | Off-Stim<br>length (hrs)<br>R/L | On-Stim<br>length (hrs)<br>R/L | Implant<br>Site | cDBS<br>Sides | Off-Stim<br>Accelerometer<br>Status | On-Stim<br>Accelerometer<br>Status |
| --- | --- | --- | --- | --- | --- | --- | --- | --- | --- | --- | --- |
| RCS02 | 54 | M | 7 | LDE 1475 | 49/5 | 135.9/73.6 | 100.2/180.0 | STN | R/L | On | On |
| RCS05 | 63 | M | 19 | LDE 955 | 45/22 | 19.1/19.2 | 32.8/24.8 | STN | R/L | On | On |
| RCS06 | 28 | F | 12 | LDE 1550 | 61/16 | 92.2/52.8 | 12.3/0 | STN | R | On | On |
| RCS07 | 40 | M | 4 | LDE 1314 | 41/14 | 94.6/108.8 | 0/0 | STN | No | On | Off |
| RCS08 | 58 | M | 12 | LDE 2100 | 44/11 | 15.0/23.3 | 136.2/101.9 | STN | R/L | On | On |
| RCS11 | 71 | M | 7 | LDE 1525 | 29/12 | 13.0/7.4 | 77.8/22.8 | STN | R/L | On | On |
| RCS12 | 58 | M | 10 | LDE 800 | 34/9 | 67.8/66.8 | 0/19.4 | STN | L | On | Off |
| RCS15 | 61 | M | 6 | LDE 570 | 35/12 | 25.4/32.3 | 12.5/8.1 | STN | R/L | On | On |
| RCS17 | 49 | M | 7 | LDE 1476 | 32/5 | 14.7/22.4 | 108.0/61.5 | STN | R/L | Off | On |
| RCS18 | 70 | M | 8 | LDE 1985 | 30/7 | 23.4/25.2 | 0/0 | STN | No | Off | Off |
| RCS20 | 76 | M | 13 | LDE 1590 | 31/10 | 27.1/26.4 | 10.5/6.6 | STN | R/L | Off | On |

Preoperative levodopa dose equivalent (LDE) and Unified Parkinson's Disease Rating Scale part III (UPDRS-III) scores are shown for each patient in the OFF- and ON-medication states. The OFF-medication state was defined as $\geq 12$ h withdrawal of antiparkinsonian medication, and the ON-medication state as 30–45 min after administration of a supratherapeutic dose of levodopa. Neural recordings were performed before (Off-Stim) and after (On-Stim) initiation of continuous subthalamic nucleus (STN) deep brain stimulation (cDBS). During recordings, the implanted pulse generator (IPG) accelerometer was either active (On) or inactive (Off). LDE, levodopa dose equivalent; STN, subthalamic nucleus.

In each hemisphere, a quadripolar paddle-type lead (10 mm intercontact spacing) was placed subdurally over the precentral and postcentral gyri, whilst a quadripolar Medtronic 3389 lead (1.5 mm intercontact spacing) was implanted in the STN as previously described^37,38^. The surgeon positioned cortical leads along a parasagittal trajectory, ensuring that two or three contacts were anterior to the central sulcus and 2–4 cm from the midline. We confirmed electrode placement intraoperatively using cone beam CT (Medtronic O-arm)^39^ fused with preoperative MRI. For each implanted hemisphere, leads were connected via 60-cm extensions (model 37087) to a Medtronic Summit RC+S interface (model B35300R) implanted in the ipsilateral pectoral region.

Precise electrode and contact locations were refined post hoc using established image analysis pipelines for deep brain stimulation^40^ and cortical electrodes^41^. Post-implantation high-resolution CT images were coregistered to preoperative T1-weighted 3 T MRI using an affine transformation^42^. Electrode placement was verified by visual inspection and brain shift correction was applied to refine subcortical anatomy coregistration when necessary^43^. Electrodes were localized based on CT artefacts, and surface projection correction was applied to align cortical leads with the MRI-derived pial surface^42^. For group analyses, electrode locations were normalised into Montreal Neurological Institute space^42^ and visualized using a Parkinson’s disease-specific subcortical atlas^44^.

### Neural data recordings

We recorded sensorimotor cortical and STN field potentials, whilst patients performed activities of daily living in their usual home environments before and after the initiation of therapeutic DBS. All recordings were obtained at least one week following surgery with patients on their usual antiparkinsonian medication. Neural signals from two bipolar channels from each cortical and STN electrode were wirelessly streamed at sampling rates of 250, 500 or 1000 Hz, from the summit RC+S interface to a Microsoft Windows-based tablet through a custom-made graphical user interface, compliant with US Food and Drug Administration code CFR 820.30 (https://openmind-consortium.github.io)^38^. During STN-targeted cDBS sessions, a single bipolar channel was streamed, configured in a “sandwich” arrangement using contacts immediately dorsal and ventral to the stimulating contact. The stimulation amplitude was increased gradually (0.1-0.2 mA every few days) to therapeutic amplitude after the stimulation onset. Patients could hold or reduce the stimulation amplitude if any adverse effects occurred. Chronic recordings were performed at a minimum of five months after the onset of stimulation^45^. **Supplementary Table 1** shows the stimulation settings for each hemisphere and patient.

### Wearable and RC+S accelerometry

During neural recording sessions, all patients wore bilateral Personal KinetiGraph (PKG) monitors (Global Kinetics Pty Ltd.) to continuously assess bradykinesia and dyskinesia severity at two-minute intervals^29,30,37,46^. PKG-derived motor scores have previously been validated against clinician rated UPDRS assessments and are widely used as objective measures of motor fluctuations in PD^37,47^. In 8 of the 11 patients, concurrent triaxial accelerometer recordings from the bilaterally implanted RC+S device^48,49^ were also obtained prior to and following the initiation of cDBS (see **Figure 1** for schematic of neural, wearable and accelerometer recordings). Accelerometer recordings were sampled at 65.12 Hz.

**Figure 1:**
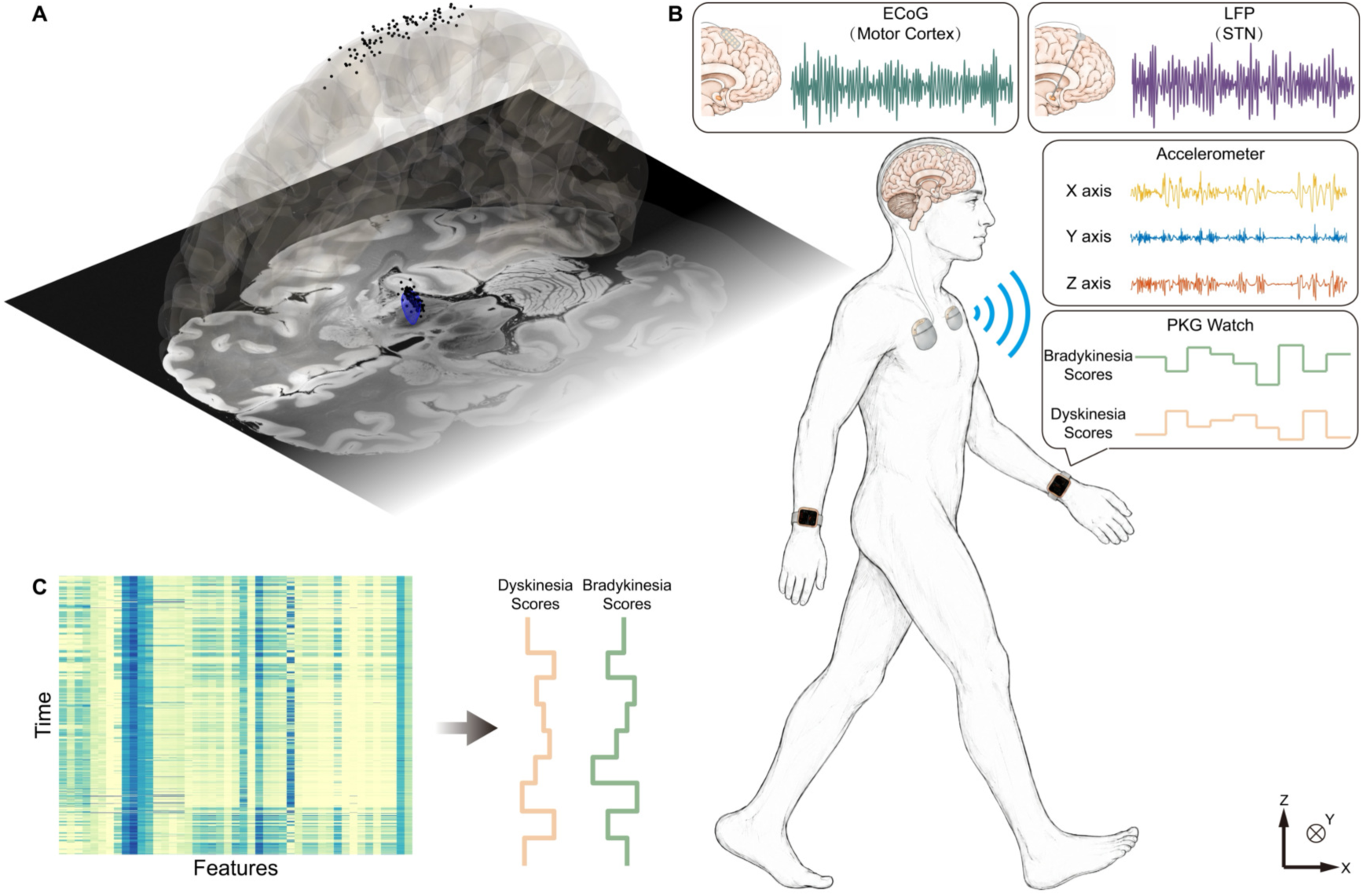
Multimodal recordings and feature extraction during naturalistic behaviour. **A**: Locations of cortical and subthalamic nucleus (STN; blue mesh) recording contacts shown in Montreal Neurological Institute (MNI) template space, overlaid on a canonical T1-weighted MRI. **B**: Simultaneous acquisition of invasive electrophysiological recordings (cortical and STN LFP), device-embedded accelerometry (RC+S), and wearable symptom estimates using the Parkinson’s KinetiGraph (PKG) during activities of daily living. **C**: Neural and accelerometer features were extracted and used to predict bradykinesia and dyskinesia scores using regression models.

### Signal processing

All RC+S data were initially processed using an open-source MATLAB toolbox to extract signal timestamps and to convert the recordings into a format compatible with MATLAB-based analysis^33^. Neural and accelerometer signals were then downsampled to 250 Hz and 64 Hz respectively, prior to a fourth-order Butterworth bandpass filter being applied; a pass band of 1–100 Hz was used for neural signals, whilst a pass band of 0.2–4 Hz was employed for RC+S accelerometer recordings to optimally capture bradykinesia and dyskinesia fluctuations as previously described^27,29^. Filtered data were segmented into non-overlapping 2-minute windows to align with PKG output intervals and facilitate downstream analysis. Neural signals from each hemisphere were paired with contralateral pectoral RC+S accelerometry data and contralateral wrist-worn PKG scores. Time stamp synchronisation was used to align the signals from the RC+S device with PKG outputs, as per previous analyses^50^. Signal segments shorter than 2 minutes at the end of each recording session were excluded.

Spectral features were computed for cortical and STN channels in each 2-minute window using a multitaper approach (frequency resolution of 1 Hz, with a taper smoothing frequency of 2.5 Hz) implemented within the FieldTrip toolbox^51^. Periodic and aperiodic components of the resulting power spectra were separated using the FieldTrip implementation of the Fitting Oscillations and One-Over-F (FOOOF) algorithm^52^. The aperiodic component was modeled as a fixed power-law to estimate an offset and an exponent (fit range: 1–100 Hz; peak-width limits: 0.5–12 Hz; maximum peaks: 3; minimum peak height: 3 dB; peak threshold: 2). Across hemispheres and stimulation conditions, the median FOOOF fit quality was high (R² = 0.93), confirming the robustness of the spectral decomposition. Mean power was then computed within PD-relevant bands—low beta (15–20 Hz), high beta (20–35 Hz), low gamma (40–70 Hz), and high gamma (70–100 Hz)—for both the periodic component and the total spectrum. Cortico-subthalamic coupling was quantified as magnitude-squared coherence within the same bands. These bands were selected based on prior links to motor symptom severity and basal ganglia–cortical dynamics in PD^6,37,53,54^.

For each 2-minute window of the preprocessed triaxial accelerometer recordings, the Euclidean norm was computed to obtain a composite measure of overall acceleration. An additional 29 movement-related acceleration features^27^ were also extracted (see **Supplementary Table 2** for further details) to capture patterns potentially indicative of motor symptoms. Feature extraction was performed independently within each 2-minute window using causal, real-time compatible operations, without incorporating information from future windows. For the ‘time below threshold’ accelerometer feature, a grid search was conducted to find the ‘optimal’ threshold leading to maximal correlation with PKG derived bradykinesia or dyskinesia scores (**Supplementary Figure 1**). All extracted neural and acceleration features were subsequently z-score normalized within hemisphere across all features to minimize inter-subject and inter-feature variability.

Each PKG score provided an estimate of motor state for the preceding 2-minute interval. To standardize interpretation, bradykinesia scores were inverted (multiplied by –1) so that higher values corresponded to greater severity, and segments with scores ≤ 0 were excluded. Periods of potential sleep were identified and removed based on sustained immobility, defined as consecutive bradykinesia scores > 80, consistent with prior work linking immobility to sleep states^12,50,55^. Segments with dyskinesia scores of 0 were also excluded to minimize the disproportionate influence of zero values on subsequent correlation analyses.

### Statistical analyses

To quantify relationships between extracted features and PKG derived bradykinesia and dyskinesia scores, we computed Spearman and Pearson correlation coefficients, as well as Kullback–Leibler (KL) divergence. Spearman correlation was primarily used to assess associations between neural and accelerometry features and PKG scores, given its robustness to outliers and sensitivity to monotonic, potentially nonlinear relationships. For comparing acceleration features with PKG scores, both Pearson correlation and KL divergence were employed to evaluate linear correspondence and distributional similarity, respectively. Based on visual inspection, an accelerometry feature was considered optimal for tracking symptom severity if it maximized linear correlation while maintaining low KL divergence with the corresponding PKG score, thereby capturing both the magnitude and distributional structure of motor state fluctuations.

All correlation analyses were conducted at the individual subject level. For each subject, correlation coefficients and associated p-values were computed using two-sided t-tests. To control for multiple comparisons and reduce the likelihood of false positives, p-values from all correlation tests were corrected using the Benjamini–Hochberg False Discovery Rate (FDR) procedure^56^. Specifically, p-values from similar feature types—such as low/high beta/gamma power, coherence, and aperiodic offset/exponent—across all hemispheres and subjects were grouped and treated as a single family of tests. These were jointly submitted to the FDR correction function to obtain the corresponding q-values, with q < 0.05 considered significant. All statistical computations were performed using MATLAB.

### Regression models

To evaluate the feasibility of predicting motor states from neural or accelerometry features, we implemented four regression models commonly applied in prior studies: Random Forest^28^, Support Vector Machine (SVM)^27^, ElasticNet, and a Fully Connected Neural Network (FNN)^57^. Implementation details and model parameters are provided in **Supplementary Table 3**. Model inputs comprised neural features, accelerometric features, or their combination, while outputs corresponded to bradykinesia or dyskinesia scores derived from the PKG monitor. Models were trained and evaluated per hemisphere, and performance was assessed using the coefficient of determination (R²). To account for imbalanced score distributions, we performed stratified 5-fold cross-validation, repeated twice. Stratification was implemented by discretizing PKG scores into eight bins to preserve their distribution across folds. Model performance was quantified as the mean R² across the resulting 10 cross-validation evaluations. To avoid data leakage, the ‘time below threshold’ accelerometry feature was optimised exclusively within the training data for each fold. Feature importance in Random Forest models was estimated using Out-of-Bag Permuted Predictor Importance^58^.

## Results

### Data Characteristics

After excluding data segments corresponding to sleep, bradykinesia scores ≤ 0, and dyskinesia scores = 0, a total of 29,568 two-minute windows were retained for bradykinesia analysis and 25,094 for dyskinesia analysis prior to DBS initiation. Similarly, for data collected during continuous DBS, we analysed 27,457 and 21,167 two-minute windows for bradykinesia and dyskinesia, respectively. For each 2-minute window, 24 neural activity and 29 device accelerometer features were extracted (see **Supplementary Table 2** for feature details), yielding feature-by-time matrices (**Figure 1**) that served as the basis for all subsequent statistical and machine learning analyses.

### Neural signals dynamically track bradykinesia and dyskinesia severity

To evaluate neural features from the STN and sensorimotor cortex as biomarkers of bradykinesia and dyskinesia prior to DBS initiation, we compared total, periodic, and aperiodic spectral power across symptom severity groups defined by wearable PKG scores^59^ (**Figure 2A**). Total power differed primarily within the cortical beta band, whereas periodic beta and aperiodic components showed more distinct modulation across severity levels. Notably, these components exhibited opposing relationships in the STN: periodic beta power increased, while aperiodic power decreased, with worsening bradykinesia. These relationships were reversed for dyskinesia.

**Figure 2:**
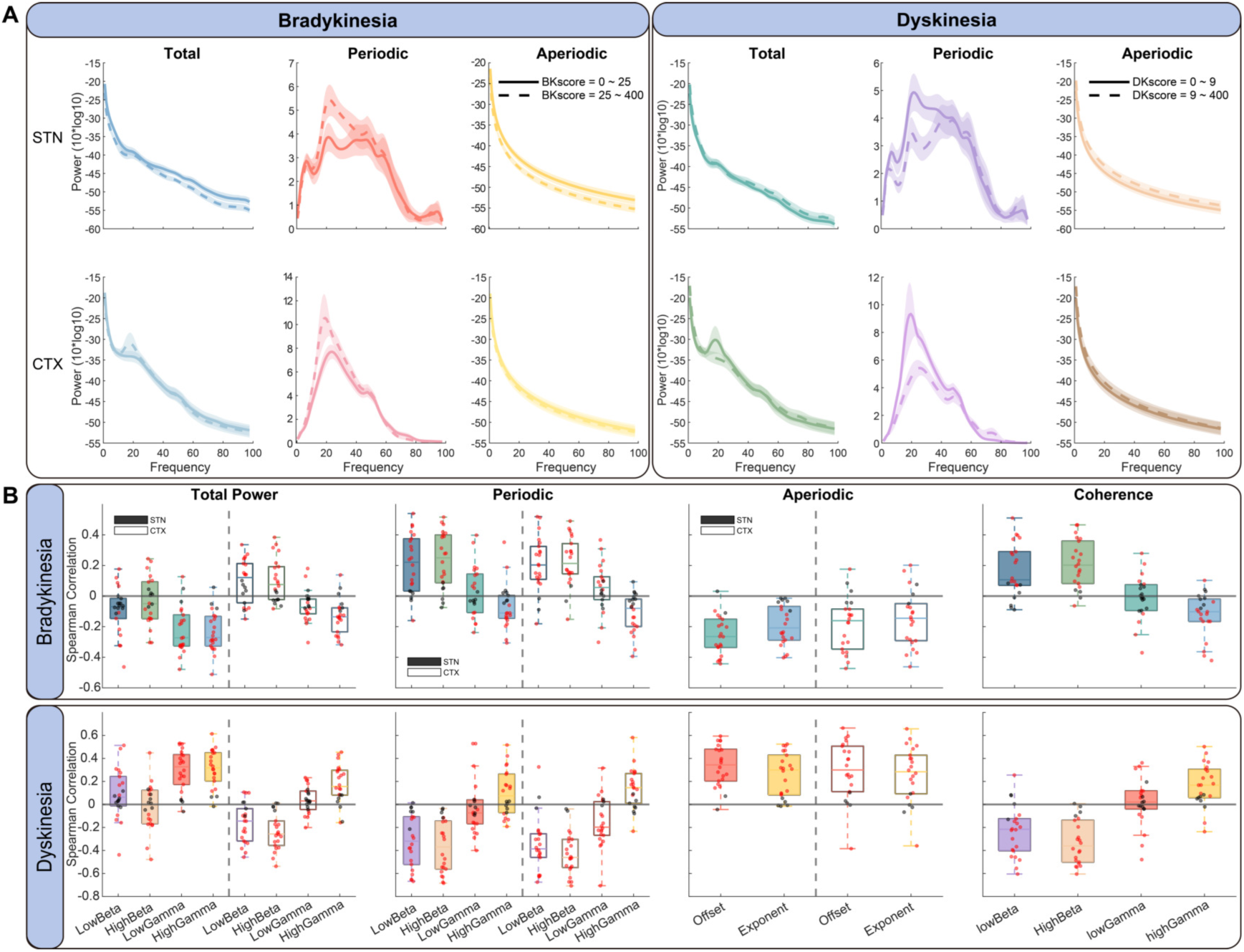
Motor cortical and STN neural signal features track Parkinsonian motor symptom severity. **A**: Power spectra for high and low symptom severity groups defined by wearable PKG scores. Lines and shaded regions represent the mean ± standard error, across hemispheres. **B**: Box-and-whisker plots showing Spearman correlations between each neural feature and bradykinesia or dyskinesia scores across hemispheres. Boxes indicate the median and interquartile range. Dark boxes represent STN features, and light boxes represent cortical (CTX) features. Different colors represent different frequency bands or aperiodic spectral fit parameters (offset or exponent). Red dots indicate hemispheres with statistically significant correlations after multiple-comparison correction (Benjamini–Hochberg false discovery rate), whereas black dots indicate non-significant associations.

To further evaluate these relationships, we computed Spearman correlations between each neural feature—including band specific total and periodic power, aperiodic offset and exponent, and cortico–STN coherence—and contralateral symptom scores. Red points in **Figure 2B** indicate hemispheres with statistically significant associations. Total STN low and high beta power showed inconsistent associations with both bradykinesia and dyskinesia across hemispheres. In contrast, periodic STN beta power robustly tracked symptom severity, increasing with worsening bradykinesia and decreasing with dyskinesia (mean Spearman coefficients for bradykinesia: total STN low beta = −0.08 ± 0.19; periodic STN low beta = 0.26 ± 0.20). For gamma band activity, total—but not periodic—power in the STN consistently scaled with motor state, increasing with improvements in bradykinesia and worsening dyskinesia.

In the cortex, both total and periodic low and high beta power increased with bradykinesia severity, whereas gamma band activity showed less consistent associations; nonetheless, high gamma power was generally anti-bradykinetic and pro-dyskinetic. For both cortical and STN sites, the aperiodic offset and exponent of the power spectrum correlated with improvements in bradykinesia. An opposite relationship was observed for dyskinesia. Finally, both low and high beta band cortico-STN coherence were predictive of worsening bradykinesia and improving dyskinesia, whereas high gamma coherence predicted improvement in bradykinesia and worsening in dyskinesia.

These findings highlight that periodic beta power is a more robust biomarker of bradykinesia than total beta power, which conflates the bradykinetic effects of periodic beta components with the prokinetic effects of the aperiodic offset. Furthermore, elevated cortico-STN beta coherence also predicted worsening bradykinesia, whereas elevations in STN gamma power, cortical gamma power, and cortico-STN gamma coherence were predictive of improvements in bradykinesia. Taken together, our results highlight a complex interplay between spatiotemporal dynamics within the cortico-STN circuit and motor fluctuations, underscoring the value of multiple spectral biomarkers for symptom tracking.

### Device accelerometry tracks bradykinesia and dyskinesia severity

To assess whether chest-implanted IPG accelerometers can track motor fluctuations, we evaluated the relationship between accelerometry features and PKG-derived bradykinesia and dyskinesia scores for each hemisphere. For this analysis, ipsilateral IPG accelerometer data were paired with PKG derived motor scores (see **Figure 1** for a schematic). Pearson correlation coefficients and Kullback–Leibler (KL) divergence were computed to quantify linear association and distributional similarity, respectively. Where necessary, feature polarity was inverted to ensure consistent directionality of correlations (**Figure 3A**).

**Figure 3:**
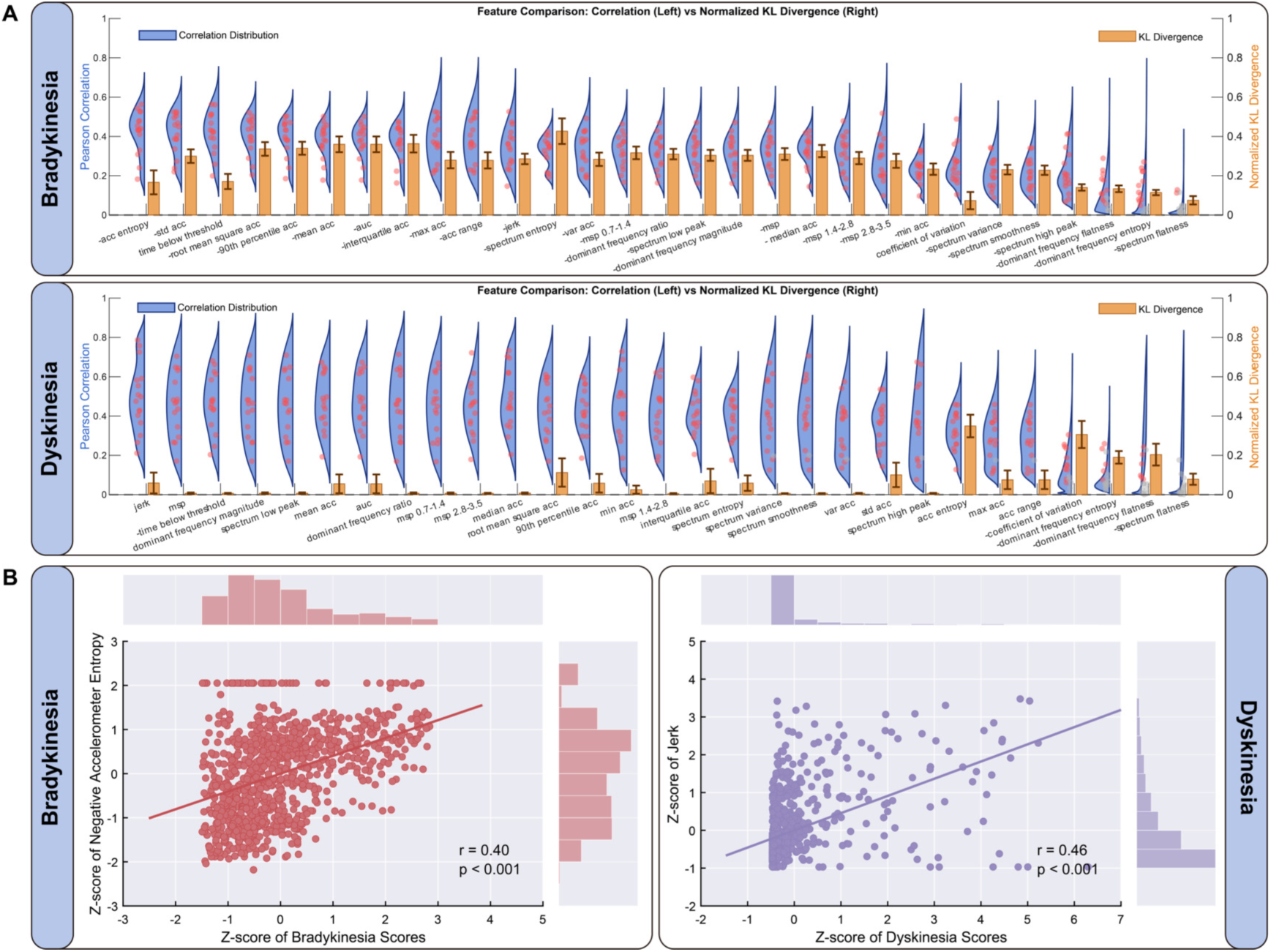
Device accelerometer features track bradykinesia and dyskinesia severity. **A:** Pearson correlation coefficients between accelerometry features and bradykinesia or dyskinesia scores across hemispheres (left axis). Blue shaded regions show the distribution of correlation coefficients for each feature. Red dots indicate hemispheres with statistically significant associations after multiple-comparison correction, whereas grey dots indicate non-significant associations. Negatively correlated features were inverted for visualisation (see main text). The right axis shows 0–1-normalised Kullback–Leibler (KL) divergence between feature and score distributions. **B:** Scatterplots with least-squares fits and marginal histograms showing associations between the optimal accelerometric feature and wearable motor scores (left: negative accelerometer entropy vs bradykinesia; right: jerk vs dyskinesia). All variables are z-scored. For visualisation, a 1-in-20 random subsample (without replacement) is shown; Pearson r and corresponding p-values were computed using all observations. Histograms show marginal distributions of features and scores.

Several accelerometry features showed significant associations with bradykinesia across hemispheres, with entropy (inverted) yielding the highest correlations alongside a low KL divergence. For dyskinesia, most features exhibited low KL divergence, with jerk showing the strongest correlations across hemispheres. Scatter plots (**Figure 3B**) illustrate these relationships between selected accelerometry features and PKG-derived scores. Together, these results demonstrate that different IPG-embedded accelerometry features robustly track bradykinesia and dyskinesia severity.

### Device accelerometry outperforms neural features for decoding motor fluctuations

Building on the observed associations between neural and accelerometry features and motor scores, we trained four regression models—Random Forest, Support Vector Machine (SVM), ElasticNet, and Fully Connected Neural Network (FNN)—to predict bradykinesia and dyskinesia severity from neural, accelerometric, or combined inputs. **Table 2** summarizes the performance of each model, expressed as the mean R^2^ under stratified cross-validation (see **Methods**) across different feature combinations. Random Forest regression achieved the highest prediction accuracy for both bradykinesia and dyskinesia, consistent with there being nonlinear structure in the data. To compare the predictive performance of different (pairwise) feature combinations within the Random Forest, we ran Wilcoxon signed-rank tests on per cross-validation fold R^2^ values obtained from the same stratified splits (**Figure 4A**). For bradykinesia prediction, accelerometer features significantly outperformed both STN (q = 5.9 × 10^-4^) and cortical activity features (q = 5.2 × 10^-6^). A similar pattern was observed for dyskinesia (accelerometry vs. STN, q = 3.0 × 10^-4^; accelerometry vs. cortical features, q = 3.9 × 10^-5^). Combining neural (STN and cortical) and accelerometry features further improved performance relative to either modality alone (**Table 2**, **Figure 4A**).

**Figure 4:**
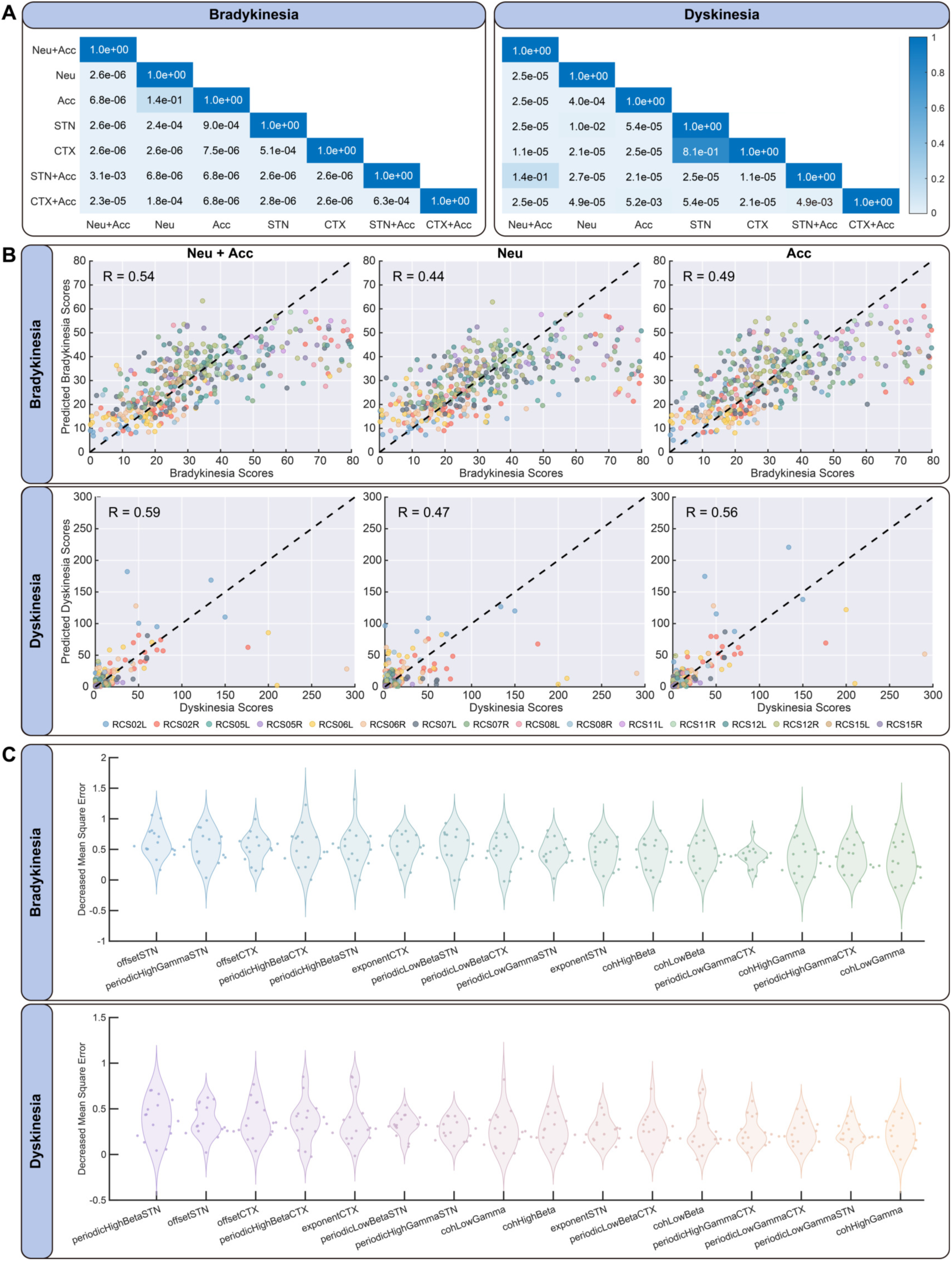
Device accelerometry outperforms neural activity features for predicting motor symptom severity. Data are shown for the Random Forest regressor. **A:** Pairwise comparisons of feature sets for bradykinesia and dyskinesia prediction. P-values were computed using Wilcoxon signed-rank tests on mean R² values across cross-validation folds. Multiple comparisons were corrected using the Benjamini–Hochberg false discovery rate, and q-values are reported. Colour indicates q-value. **B:** Predicted versus observed bradykinesia and dyskinesia scores across hemispheres for models trained using neural features (Neu), accelerometry features (Acc), or their combination (Neu+Acc). Points lie on the diagonal (dashed line) for perfect predictions. For visualisation, a 1-in-50 random subsample of validation data is shown; R² values were computed using all observations. **C:** Feature importance for bradykinesia and dyskinesia prediction. Neural features are ordered by mean importance across hemispheres, quantified using out-of-bag (OOB) permutation importance. Each point represents one hemisphere.

**Table 2:** Predictive performance of four regressors for wearable bradykinesia and dyskinesia scores using neural, accelerometric, and combined features. Data are shown for recordings obtained without (cDBS Off) and during continuous DBS (cDBS On) of the STN.

| Stimulation | Behavior | Regressor | Neural +<br>Acc (R2) | Neural<br>(R2) | Acc<br>(R2) | STN<br>(R2) | CTX<br>(R2) | STN +<br>Acc (R2) | CTX +<br>Acc (R2) |
| --- | --- | --- | --- | --- | --- | --- | --- | --- | --- |
| cDBS Off | Bradykinesia | **Random Forest** | **0.2925** | **0.1951** | **0.2430** | **0.1713** | **0.1436** | **0.2857** | **0.2724** |
|  |  | SVM | 0.1180 | 0.1137 | 0.1659 | 0.1242 | 0.1067 | 0.1559 | 0.1465 |
|  |  | Elastic Net | -2.9714 | 0.1361 | -2.2758 | 0.0865 | 0.0234 | -1.3605 | -2.7188 |
|  |  | Fully Connected<br>Neural Network | 0.2455 | 0.1143 | 0.2180 | 0.0803 | 0.0231 | 0.2409 | 0.2321 |
|  | Dyskinesia | **Random Forest** | **0.3463** | **0.2252** | **0.3099** | **0.1934** | **0.1807** | **0.3428** | **0.3258** |
|  |  | SVM | 0.0022 | 0.0842 | 0.0593 | 0.0617 | 0.0771 | 0.0265 | 0.0221 |
|  |  | Elastic Net | -0.0265 | -0.5189 | -0.8178 | 0.0860 | 0.0600 | -0.4174 | -0.1580 |
|  |  | Fully Connected<br>Neural Network | 0.2468 | 0.1472 | 0.2583 | 0.1546 | 0.1087 | 0.2124 | 0.2171 |
| cDBS On | Bradykinesia | **Random Forest** | **0.2795** | **0.2154** | **0.2425** | **0.1105** | **0.1806** | **0.2649** | **0.2660** |
|  |  | SVM | 0.0687 | 0.0898 | 0.1523 | 0.0482 | 0.1347 | 0.1197 | 0.1271 |
|  |  | Elastic Net | 0.2143 | 0.1506 | 0.1005 | 0.0410 | 0.1355 | 0.2190 | 0.1842 |
|  |  | Fully Connected<br>Neural Network | 0.2219 | 0.1056 | 0.2326 | 0.0079 | 0.0660 | 0.2313 | 0.2351 |
|  | Dyskinesia | **Random Forest** | **0.3433** | **0.2521** | **0.3149** | **0.1348** | **0.2153** | **0.3352** | **0.3384** |
|  |  | SVM | -0.0622 | 0.0097 | 0.0275 | 0.0211 | 0.0689 | -0.0341 | -0.0382 |
|  |  | Elastic Net | 0.2988 | 0.1665 | 0.3012 | 0.0624 | 0.1425 | 0.2951 | 0.2824 |
|  |  | Fully Connected<br>Neural Network | 0.2862 | 0.1200 | 0.3005 | 0.0385 | 0.1620 | 0.3012 | 0.2835 |

**Feature sets:** Neural+Acc; Neural only (STN+CTX); Acc only; STN only; CTX only; STN+Acc; CTX+Acc.
**Performance metric:** mean $R^2$ (coefficient of determination) from stratified 5-fold cross-validation repeated twice (10 folds total). Negative $R^2$ values indicate model performance worse than a mean predictor and arise from cross-validated evaluation.
**Abbreviations:** Neural = neural features (including both STN and cortical features); Acc = accelerometric features; STN = subthalamic nucleus features; CTX = cortical features; SVM = support vector machine.

These results demonstrate that device-embedded accelerometry provides a robust signal for decoding Parkinsonian motor states. Notably, a feature set combining only STN and accelerometry features approached the performance of the full model (R² for device accelerometry and STN features = 0.28; R² for device accelerometry, cortical and STN features = 0.29), suggesting a clinically practical configuration for adaptive DBS in settings where cortical recordings are unavailable.

To further assess model performance, we compared predicted and observed wearable scores (**Figure 4B**, **Supplementary Figure 2**). Predictions closely tracked true values across a broad range of symptom severity, with increased dispersion at higher scores (bradykinesia > 50, dyskinesia > 150). Importantly, models captured the full dynamic range of the data, indicating that performance was not driven by outliers.

Feature importance was estimated using out-of-bag permutation importance (**Figure 4C**), ranked by the mean decrease in out-of-bag mean squared error across hemispheres. For bradykinesia, the most informative features included periodic high gamma power in the STN, as well as periodic high beta power and aperiodic offset in both STN and cortex. For dyskinesia, key features included periodic high beta power and aperiodic offset in both cortex and STN, as well as the cortical aperiodic exponent. These findings parallel the correlation analyses and highlight the importance of both periodic beta and aperiodic components in tracking symptom severity.

### Correlations between neural signals and motor symptoms are recapitulated using accelerometry-derived features

Using the optimal accelerometry features identified above as surrogate markers of bradykinesia and dyskinesia, we computed Spearman correlations between these features and neural signals across hemispheres. The resulting patterns closely matched those observed between neural features and PKG-derived scores (**Figure 5**).

**Figure 5:**
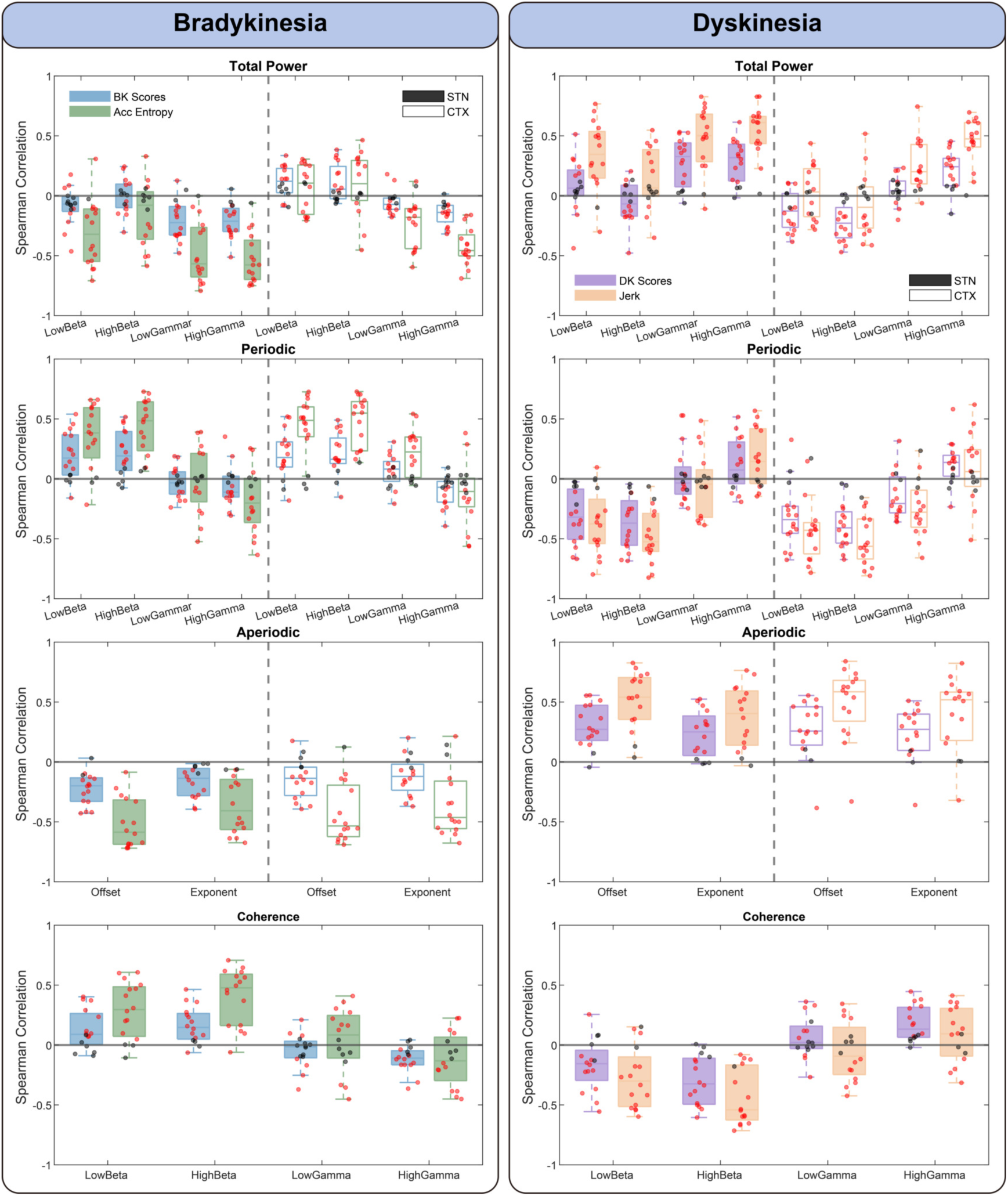
Optimal accelerometer features recapitulate neural correlates of motor symptom severity. Hemisphere-wise Spearman correlations between neural features and either PKG-derived motor scores or the corresponding optimal accelerometry-derived surrogate. Colours within each subplot denote correlation type (PKG score vs. device accelerometry; see legend). Dark shaded panels indicate STN features, whereas light shaded panels indicate motor cortical (CTX) features. Red points indicate hemispheres with statistically significant correlations after Benjamini–Hochberg false discovery rate correction (q < 0.05), whereas black points indicate non-significant associations. Offset, aperiodic offset; exponent, aperiodic exponent. STN, subthalamic nucleus; CTX, motor cortex.

Accelerometry-based correlations were generally stronger and more frequently significant across hemispheres than those obtained using wearable scores. For most neural metrics (excluding total power), accelerometry also improved cross-hemisphere consistency, reflected by tighter clustering of significant effects and few non-significant correlations. These improvements were most pronounced for aperiodic parameters and beta-band measures and were weaker in the gamma band. Together with earlier results (**Figure 3**), our findings demonstrate that device-embedded accelerometry provides a robust surrogate for motor symptom severity and preserves the underlying relationships between neural activity and motor state within the cortico–STN network.

### Neural and accelerometry biomarkers are differentially impacted by continuous STN stimulation

To assess the stability of neural and accelerometry-derived biomarkers during continuous STN DBS (cDBS), we repeated the above analyses using recordings acquired under stimulation. Under cDBS, most hemispheres retained significant associations with symptom severity; however, both the strength and cross-hemisphere consistency of neural–symptom relationships were markedly reduced. This attenuation was particularly evident for STN periodic beta power and cortical aperiodic components, reflected by reduced Spearman correlation magnitudes and an increased prevalence of opposing correlation signs across hemispheres (**Figure 6A**). In contrast, STN aperiodic parameters, cortical periodic power, and cortico–STN coherence preserved comparatively stable associations with PKG scores under stimulation (**Figure 6A**). These findings suggest that stimulation-induced modulation of oscillatory activity—particularly beta-band dynamics—decouples several canonical spectral biomarkers from motor state, whereas aperiodic and network-level measures remain informative.

**Figure 6:**
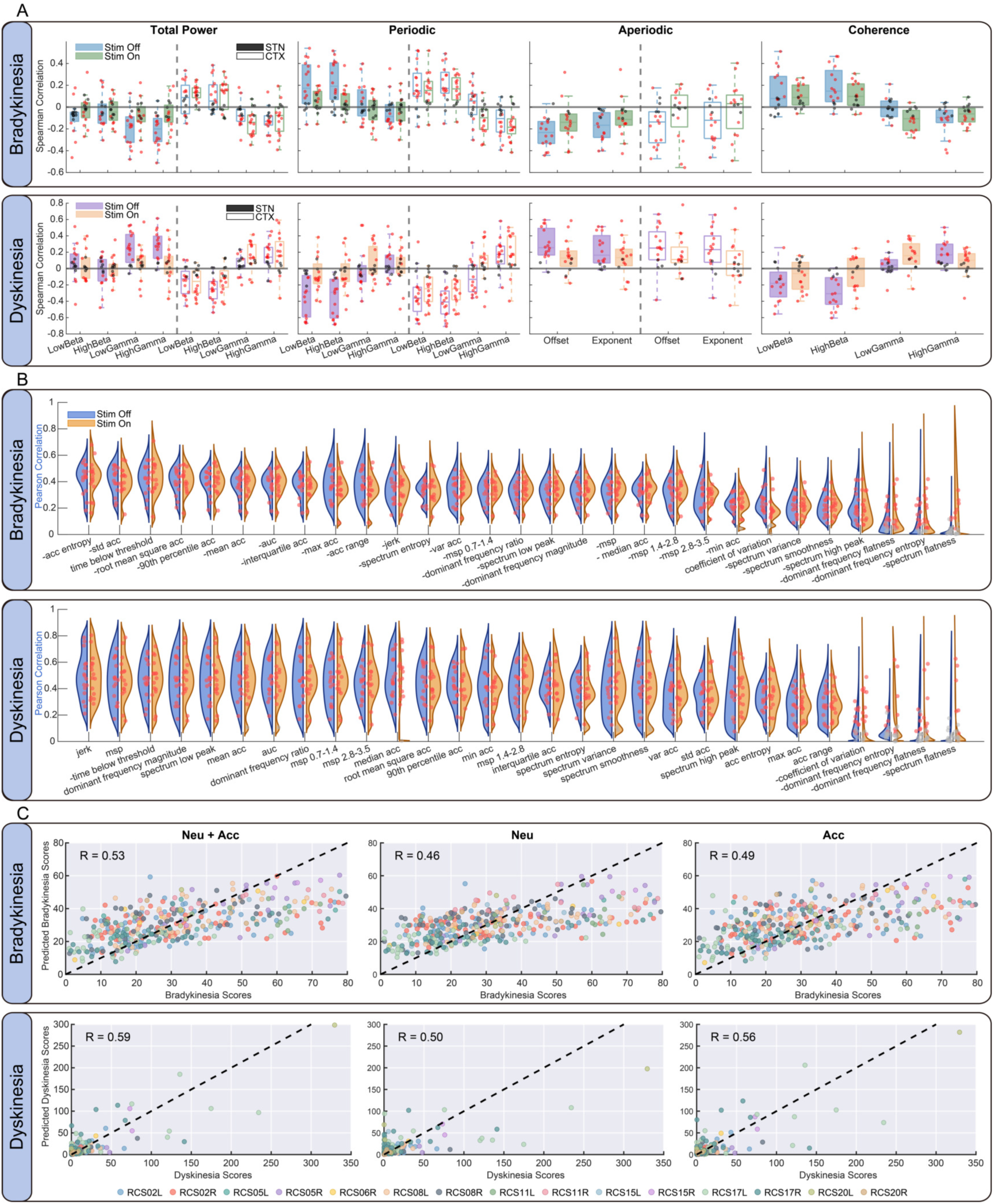
Comparison of neural and accelerometer features for tracking motor symptoms during continuous STN DBS. **A:** Spearman correlations between neural features and bradykinesia or dyskinesia scores, shown separately for the stimulation off and on conditions across hemispheres. Boxes indicate the median and interquartile range. Dark boxes represent STN features, and light boxes represent cortical (CTX) features. Colours denote frequency bands or aperiodic parameters (offset and exponent). Red points indicate hemispheres with statistically significant correlations after Benjamini–Hochberg false discovery rate correction, whereas black points indicate non-significant associations. **B:** Pearson correlations between accelerometry features and bradykinesia or dyskinesia scores across hemispheres, shown for the stimulation off and on conditions. Red points indicate statistically significant correlations after multiple-comparison correction, whereas grey points indicate non-significant associations. Negatively correlated features were inverted for visualisation. **C:** Predicted versus observed bradykinesia and dyskinesia scores under continuous stimulation, shown for models using neural features (Neu), accelerometry features (Acc), or their combination (Neu+Acc). For visualisation, a 1-in-50 random subsample of validation data from each cross-validation fold is shown. The coefficient of determination (R²) was computed using all observations. Points on the diagonal (dashed line) indicate perfect predictions.

By comparison, accelerometry-derived features exhibited minimal degradation under cDBS. Correlations between accelerometry features and wearable symptom scores were largely preserved (**Figure 6B**), and the optimal accelerometry feature set maintained expected relationships with neural activity (**Supplementary Figure 3**). Consistent with this stability, accelerometry-only models outperformed neural activity models for both bradykinesia and dyskinesia prediction under stimulation (**Table 2** and **Supplementary Figure 4**), highlighting the robustness of kinematic features for symptom tracking. For bradykinesia prediction, accelerometry significantly outperformed both STN (q = 5.5 × 10^-6^) and cortical features (q = 9.9 × 10^-6^). A similar pattern was observed for dyskinesia (accelerometry vs STN, q = 2.7 × 10^-5^; accelerometry vs cortex, q = 1.4 × 10^-4^). Combining STN and cortical features improved neural model performance. However, accelerometry-based models continued to outperform combined neural models for both bradykinesia (q = 7.6 × 10^-3^) and dyskinesia (q = 4.6 × 10^-4^).

Regression analyses further demonstrated effects of stimulation on decoding performance. Bradykinesia prediction using both neural and accelerometry features declined modestly under cDBS (R² off stimulation: 0.292; R² cDBS: 0.247), whereas dyskinesia prediction remained stable (**Figure 6C**; **Table 2**). STN spectral features showed reduced contributions to model performance during stimulation, consistent with their weakened correlations. These findings indicate that accelerometry can compensate for stimulation-induced degradation of neural biomarkers.

Feature importance analyses supported these observations. Following stimulation onset, the contribution of cortical features increased, whereas the importance of STN periodic components—particularly beta power—declined substantially (**Supplementary Figure 4**). In contrast, STN aperiodic parameters remained among the most informative predictors, reinforcing the relative stability of non-oscillatory spectral components under cDBS. Additionally, cortical gamma-band activity increased in importance during stimulation, suggesting that stimulation-associated changes in cortical synchrony may provide complementary information for tracking motor state^45,46,60^.

## Discussion

This study demonstrates that device-embedded accelerometry can accurately track motor symptom fluctuations in Parkinson’s disease, providing a practical and readily deployable signal for closed-loop control. Using an investigational DBS system capable of streaming motor cortical, STN, and accelerometry signals during naturalistic behaviour, alongside high-temporal-resolution PKG measurements, we analysed over 1,900 hours of neural recordings for bradykinesia and 1,500 hours for dyskinesia across both off- and on-stimulation states. We identify distinct neural and accelerometry-derived features that optimally track motor symptom severity and, to our knowledge, provide the first demonstration that implant-based accelerometry and neural recordings can jointly track Parkinsonian motor states during naturalistic behaviour and remain robust under STN stimulation.

These results also provide insight into the relationship between neural biomarkers and motor symptoms, revealing distinct contributions of oscillatory (periodic) and non-oscillatory (aperiodic) spectral components. Off stimulation, periodic beta power and cortico–STN coherence tracked worsening bradykinesia; however, total STN beta power—the most widely used biomarker for aDBS^17,61^—was less reliable, as it conflates opposing contributions from prokinetic aperiodic offset and bradykinetic periodic beta activity. Under stimulation, suppression of oscillatory activity further weakened the association between periodic beta power and motor symptoms. In contrast, STN aperiodic parameters, cortical periodic power, and cortico–STN coherence retained comparatively stable relationships with symptom severity, indicating that non-oscillatory and network-level features provide more robust biomarkers when oscillatory peaks are attenuated. These findings support the use of aperiodic components, rather than total beta power, for clinical aDBS control. They further demonstrate that combining STN and cortical features can improve motor state decoding, particularly in cases where pathological STN beta activity is weak or absent. This may be especially relevant for a substantial subset of patients (>15% of hemispheres) who do not exhibit elevated STN beta activity, or in whom beta oscillations are heavily suppressed by stimulation^61,62^.

A key translational finding is that device accelerometry consistently outperformed neural features for predicting motor symptom severity across both stimulation states. Although combining STN and cortical signals improved neural model performance, accelerometry retained superior predictive accuracy, particularly for dyskinesia. Given that cortical recordings are not routinely available in clinical DBS systems, the combination of STN signals and device-embedded accelerometry represents a practical and scalable configuration for closed-loop aDBS.

Accelerometry also offers several practical advantages over neural recordings that may favour its use in device deployment. First, accelerometry is less susceptible to physiological and stimulation-related artefacts (e.g. ECG^63,64^ or stimulation interference^65^). Second, accelerometers rely on low-power MEMS (microelectromechanical systems) sensors, reducing hardware complexity and energy requirements compared with chronic high-fidelity neural sensing. Together, these properties position accelerometry as a robust and clinically viable signal for real-time symptom tracking.

Taken together, our findings support a multimodal approach to aDBS, in which accelerometry provides a stable behavioural readout that complements neural biomarkers, particularly under conditions where stimulation degrades neural signal fidelity. We next consider these findings in the context of prior work and discuss study-specific limitations.

### Distinct contributions of neural activity components to motor states

Our findings support the view that periodic beta activity within the motor cortical-STN circuit exerts bradykinetic effects^14,66^, whereas narrowband gamma band activity is generally prokinetic but also associated with dyskinesia^37^. Optimal motor function may therefore depend on suppressing pathological beta oscillations while maintaining gamma activity within a physiological range that supports movement without triggering dyskinesia^67^.

Consistent with this framework, periodic high-beta STN power was a strong predictor of bradykinesia across hemispheres off stimulation, supporting a pathological role for cortico–STN transmission via the hyperdirect pathway^18^. Under stimulation, however, the importance of STN beta activity declined, consistent with suppression of oscillatory synchrony in this frequency range^13,21,68,69^. In contrast, cortical periodic power—particularly in the gamma band—became a more prominent predictor, suggesting that stimulation-induced changes in cortical gamma synchrony may provide a complementary, stimulation-resilient marker of motor state^60^.

Importantly, aperiodic cortical and STN neural activity components (both the aperiodic offset and exponent) were consistently prokinetic. These observations corroborate previous investigations of the role of aperiodic neural activity within the STN^20,21^. Mechanistically, periodic and aperiodic components are thought to reflect separable processes^20,52,70^: periodic oscillations index synchronous neuronal bursting^20,53,71,72^, whereas broadband aperiodic activity reflects neural spiking activity that occurs outside of bursts^20,73–75^. The persistence of aperiodic–symptom relationships under stimulation suggests that these features capture aspects of neural activity that are not fully suppressed by stimulation-driven reductions in oscillatory synchrony^20,21^.

This interpretation is consistent with classical basal ganglia models, in which elevated STN spiking contributes to Parkinsonian motor impairment^70,76,77^. In this framework, reductions in aperiodic exponent may reflect increased spiking arising from diminished pallidal inhibition, whereas increases in the exponent may index restoration of inhibitory balance^21^. Similarly, the aperiodic offset is closely linked to population firing rates^74,75^ and is strongly correlated with the exponent within the STN^78^.

Our results also highlight differential dominance of periodic and aperiodic spectral components within the STN and cortex (**Figure 2**). Specifically, aperiodic components appear to be the dominant signals in the STN, likely reflecting exaggerated spiking activity^20–22,74,75^. In contrast, periodic beta oscillations tend to be more pronounced in the cortex^50^ and are recognised to trigger pathological synchronisation of STN neuronal firing^79–81^.

Together, these findings extend current aDBS approaches, which typically rely on narrow-band beta power and clinician-defined thresholds^16,17^. Decomposing neural spectra into periodic and aperiodic components provides a richer and more stable representation of motor state, with potential to improve the robustness and adaptability of closed-loop stimulation strategies.

### Device accelerometer features reliably track bradykinesia and dyskinesia

Bradykinesia and dyskinesia reflect opposing states of dopaminergic tone in PD^82,83^. Prior studies have shown that trunk-mounted sensors can capture both motor states^32,84–87^, motivating the use of the embedded accelerometer within the RC+S device. Across a range of features, accelerometry showed strong relationships with PKG-derived motor scores. For bradykinesia, entropy of acceleration magnitude was the most informative feature, consistent with reduced movement velocity and complexity^1^. In contrast, dyskinesia was best captured by jerk, reflecting the rapid and irregular acceleration changes characteristic of hyperkinetic movements^2,32^. Importantly, these relationships were largely preserved under cDBS, indicating that implant-based accelerometry provides a stimulation-resilient signal that can complement neural biomarkers.

### Implications for closed-loop DBS and study limitations

The stimulation-dependent decoupling of several STN periodic biomarkers from symptoms has direct implications for aDBS design. First, it suggests that controllers trained exclusively off stimulation may not generalise optimally once stimulation is active, motivating stimulation-state–aware decoding (e.g., separate models for off vs on stimulation, or adaptive weighting of features as a function of stimulation state)^88–90^. Second, the relative stability of accelerometry, STN aperiodic activity, and cortico–STN coherence under cDBS supports a hierarchical control framework in which accelerometry provides a robust estimate of clinical state, while neural features contribute mechanistic specificity and enhance responsiveness during behavioural transitions or in contexts where kinematic signals are ambiguous^91,92^. Finally, the increased importance of cortical periodic gamma under stimulation raises testable hypotheses about stimulation-driven cortical network dynamics and suggests that cortical sensing—where available—may offer complementary, stimulation-resilient biomarkers for future bidirectional DBS systems^92,93^.

Our findings should be interpreted in light of several important limitations. Firstly, wrist-worn PKG devices may not accurately capture axial or lower limb motor symptoms^94–96^, potentially reducing the magnitude of our observed relationships between neural spectra and symptom scores. In contrast, chest-implanted accelerometers may be better positioned to capture both axial and upper-limb movement, although we were not able to apply proprietary PKG algorithms to derive directly comparable symptom scores from IPG signals^46^. Secondly, accelerometry-based decoding may, in part, reflect direct measurement of movement rather than a latent clinical state per se. As wearable PKG scores are themselves derived from kinematic features, the superior performance of accelerometry relative to neural signals may partly arise from shared measurement domains. Nevertheless, PKG-derived metrics correlate strongly with clinician-rated motor severity, including UPDRS scores, and are widely used as objective proxies of clinical state in Parkinson’s disease. In this context, accelerometry may capture behaviourally expressed motor impairment with established clinical relevance, rather than merely raw movement^47,97^.

While this distinction is unlikely to limit the utility of accelerometry for closed-loop control —where behavioural readouts may be the most relevant target^91^ — it does highlight an important conceptual challenge in interpreting decoding performance. Our models likely capture a composite of motor state and movement expression, rather than fully dissociating underlying dopaminergic or clinical state from behaviour.

More broadly, although this study establishes proof-of-principle that device-embedded accelerometry can provide a robust, stimulation-resilient signal for motor state tracking, prospective clinical trials will be required to determine whether integrating accelerometry with neural recordings improves the efficacy and side-effect profile of aDBS. Such studies will be critical to defining the optimal balance between behavioural and neural control signals in closed-loop neuromodulation systems.

## Data Availability

De-identified, processed neural and accelerometry data as well as wearable data can be provided upon request according to the data-sharing policies of the National Institutes of Health (NIH).

## Code Availability

MATLAB code and example data for reproducing analysis and figures are deposited on the GitHub repository: TonyTaoLiu/Device-Accelerometry

## Appendix

**Supplementary Table 1:** Continuous DBS parameters used for each patient and hemisphere.

| Subject | Hemisphere | Amplitudes (mA) | pulse width (us) | Frequency (Hz) | Stim Contact |
| --- | --- | --- | --- | --- | --- |
| RCS02 | Left | 2.4 - 2.6 | 60 | 130.2 | c+2- |
|  | Right | 3.1 - 3.5 | 60 | 130.2 | c+1- |
| RCS05 | Left | 1.8 - 3.0 | 60 | 130.2 | c+1- |
|  | Right | 1.3 - 1.8 | 60 | 130.2 | c+1- |
| RCS06 | Left | 1.2 | 60 | 158.7 | c+2- |
|  | Right | 1.2 | 60 | 166.7 | c+1- |
| RCS08 | Left | 2.7 - 4.7 | 60 | 130.2 | c+2- |
|  | Right | 1.0 - 3.3 | 60 | 130.2 | c+2- |
| RCS11 | Left | 2.3 - 3.5 | 60 | 130.2 | c+2- |
|  | Right | 1.5 - 2 | 60 | 130.2 | c+2- |
| RCS12 | Left | 1.7 - 3.2 | 60 | 130.2 | c+2- |
| RCS15 | Left | 3.3 - 3.5 | 60 | 130.2 - 150.6 | c+1- |
|  | Right | 3.8 - 4.4 | 70 | 130.2 - 150.6 | c+1- |
| RCS17 | Left | 2.3 - 3 | 60 | 130.2 | c+2- |
|  | Right | 1.6 - 2.5 | 60 | 129.9 - 130.2 | c+2- |
| RCS20 | Left | 1.2 - 3.3 | 60 | 130.2 | c+1- |
|  | Right | 1.2 - 2.1 | 60 | 130.2 | c+1- |

**Supplementary Table 2:**
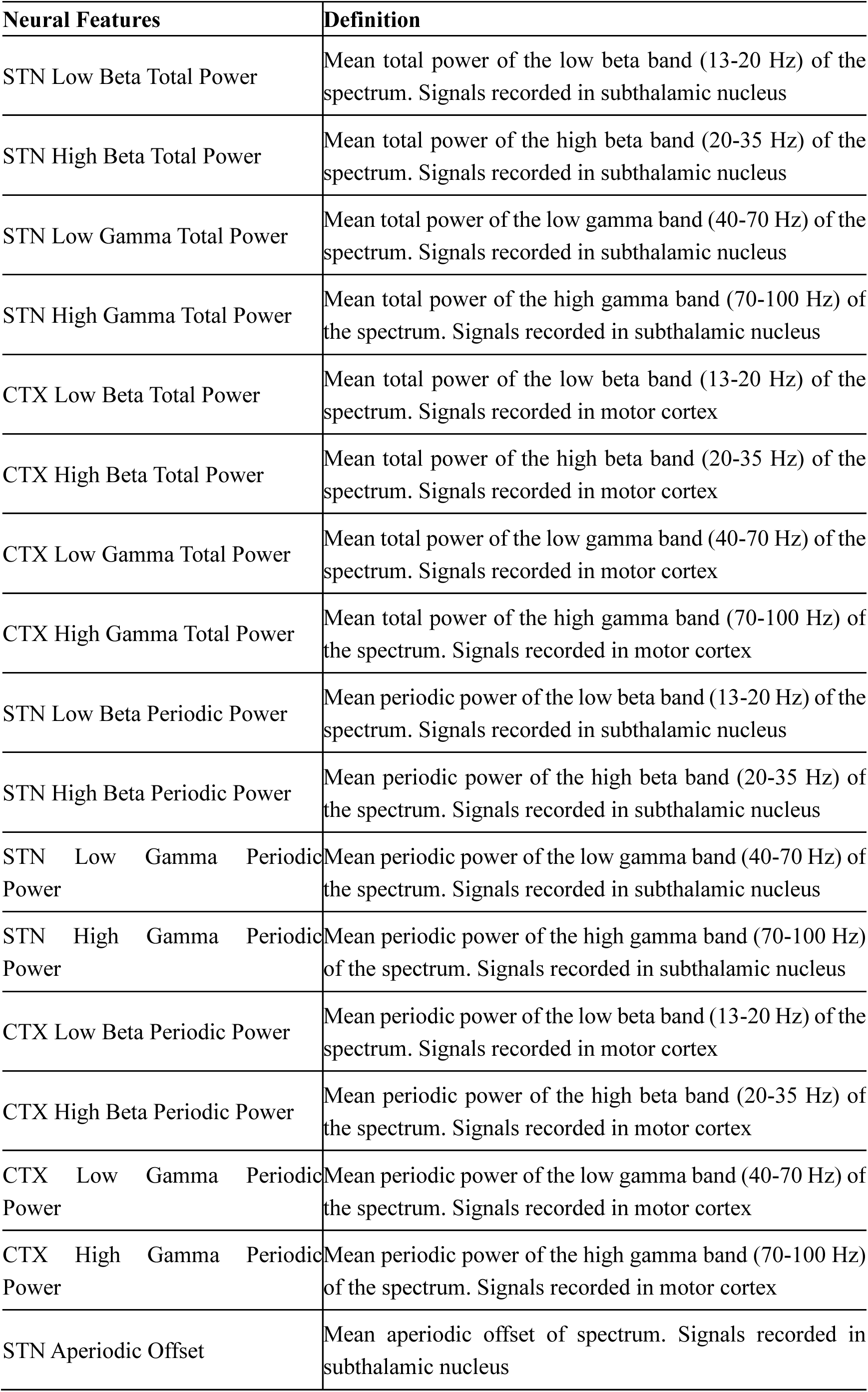
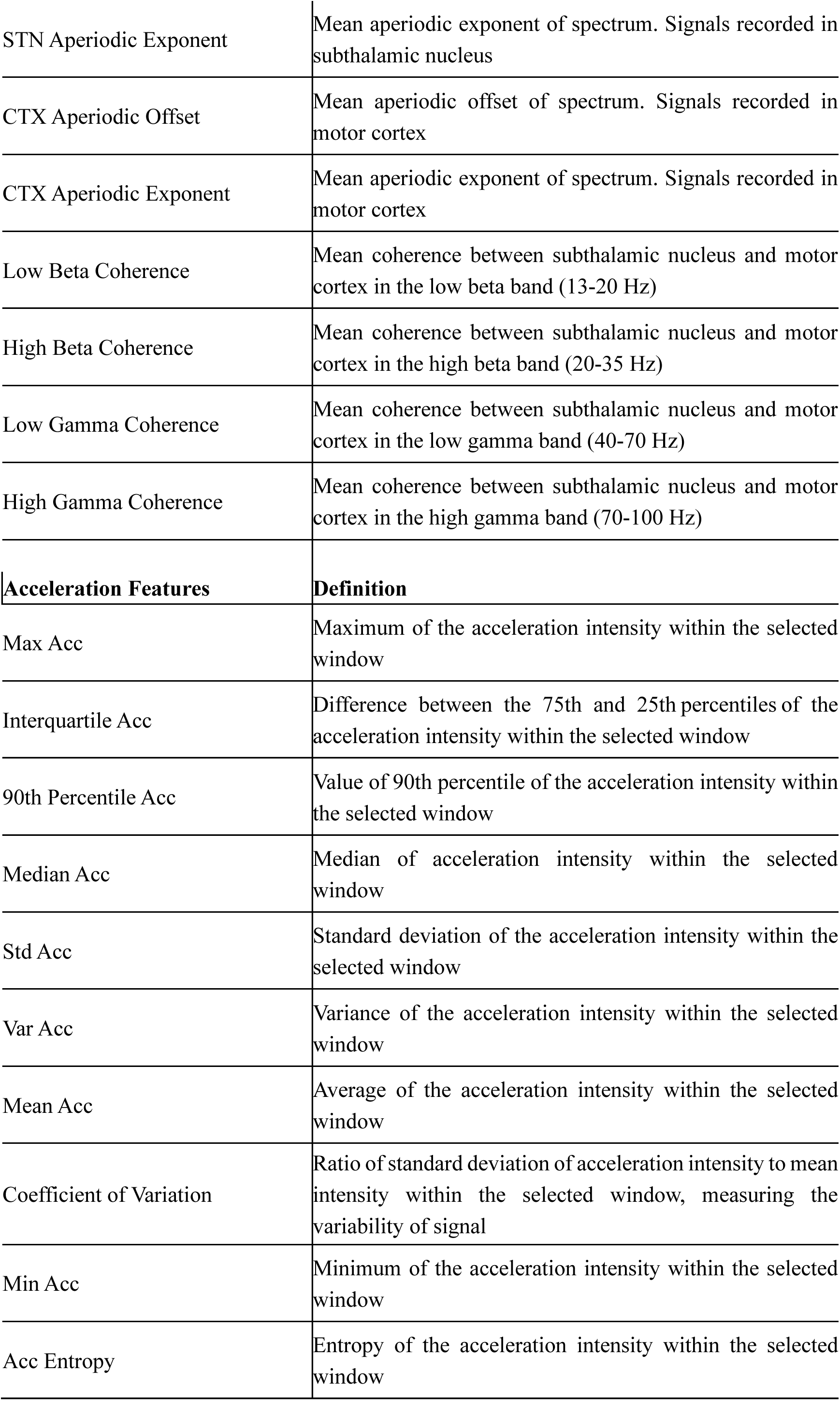
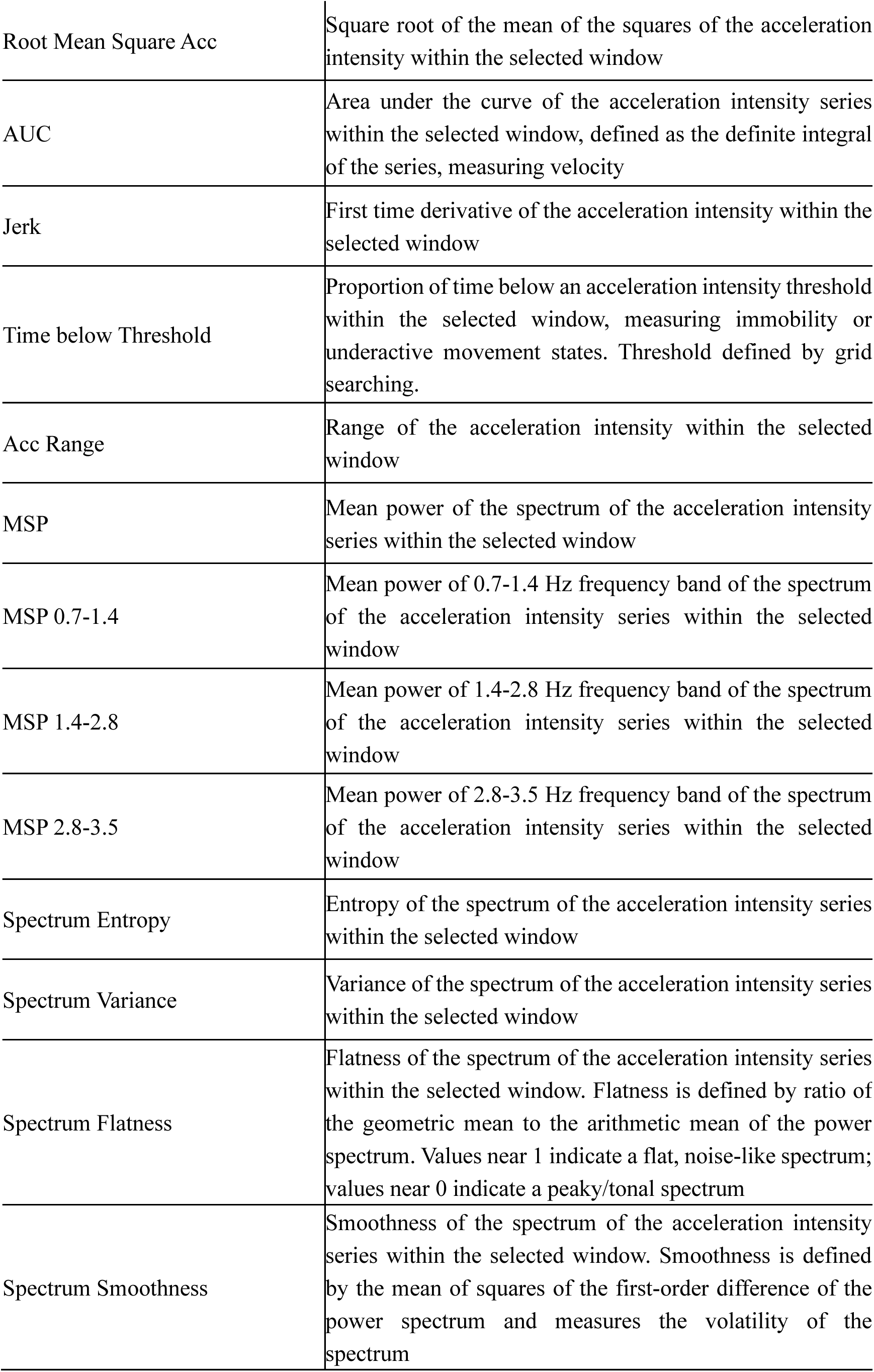
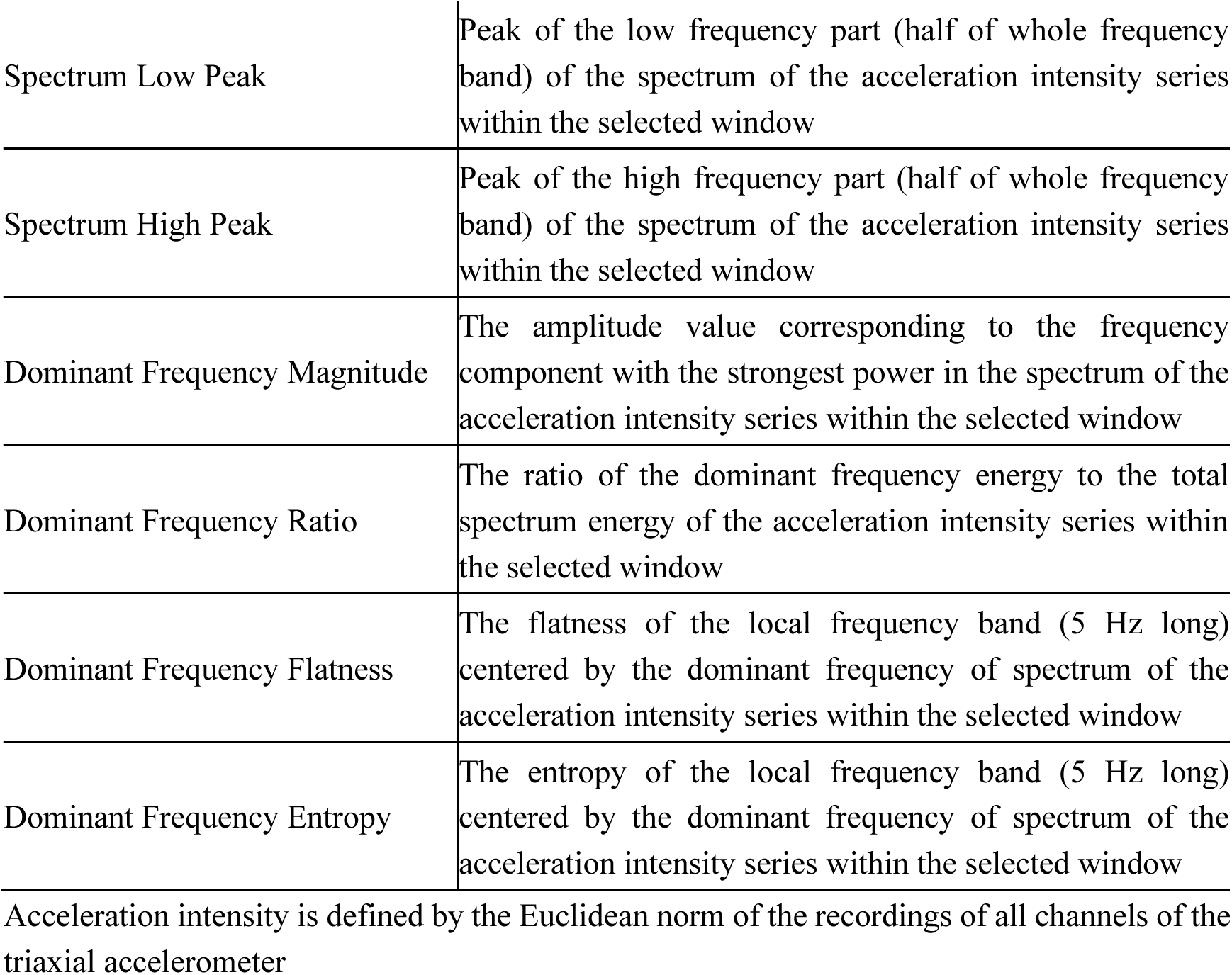
Feature Definitions Acceleration intensity is defined by the Euclidean norm of the recordings of all channels of the triaxial accelerometer.

**Supplementary Table 3:** Parameters for Regression Models All models were implemented in MATLAB 2024b. D in featureInputLayer(D) refers to the dimension of input features.

| Model | MATLAB Function | Parameters |
| --- | --- | --- |
| Random Forest | TreeBagger | NumTrees', 200<br>'Method', 'regression'<br>'NumPredictorsToSample', Number of predictors/3<br>'MinLeafSize', 5<br>'SampleWithReplacement', 'on'<br>'OOBPredictorImportance', 'on' |
| SVM | fitrsvm | KernelFunction', 'rbf'<br>'KernelScale', 'auto'<br>'BoxConstraint', 1<br>'Epsilon', 0.1<br>'Standardize', 'false'<br>'Solver', 'SMO' |
| Elastic Net | lasso | Alpha', 0.5<br>'CV', 10<br>'Lambda', selected via 10-fold cross-validation (CV = 10) to minimize the mean squared error.<br>'Standardize', 'true'<br>'RelTol', 1e4<br>'NumLambda', 100<br>'MaxIter', 1e4 |
| Fully Connected Network | trainNetwork | Network Structure: featureInputLayer(D)-fullyConnectedLayer(64)-reluLayer-fullyConnectedLayer(32)-reluLayer-fullyConnectedLayer(1)-regressionLayer<br>Loss function: Mean Square Error<br>'Optimizer', 'adam'<br>'MaxEpochs', '50'<br>'MiniBatchSize', 32 |

**Supplementary Figure 1:**
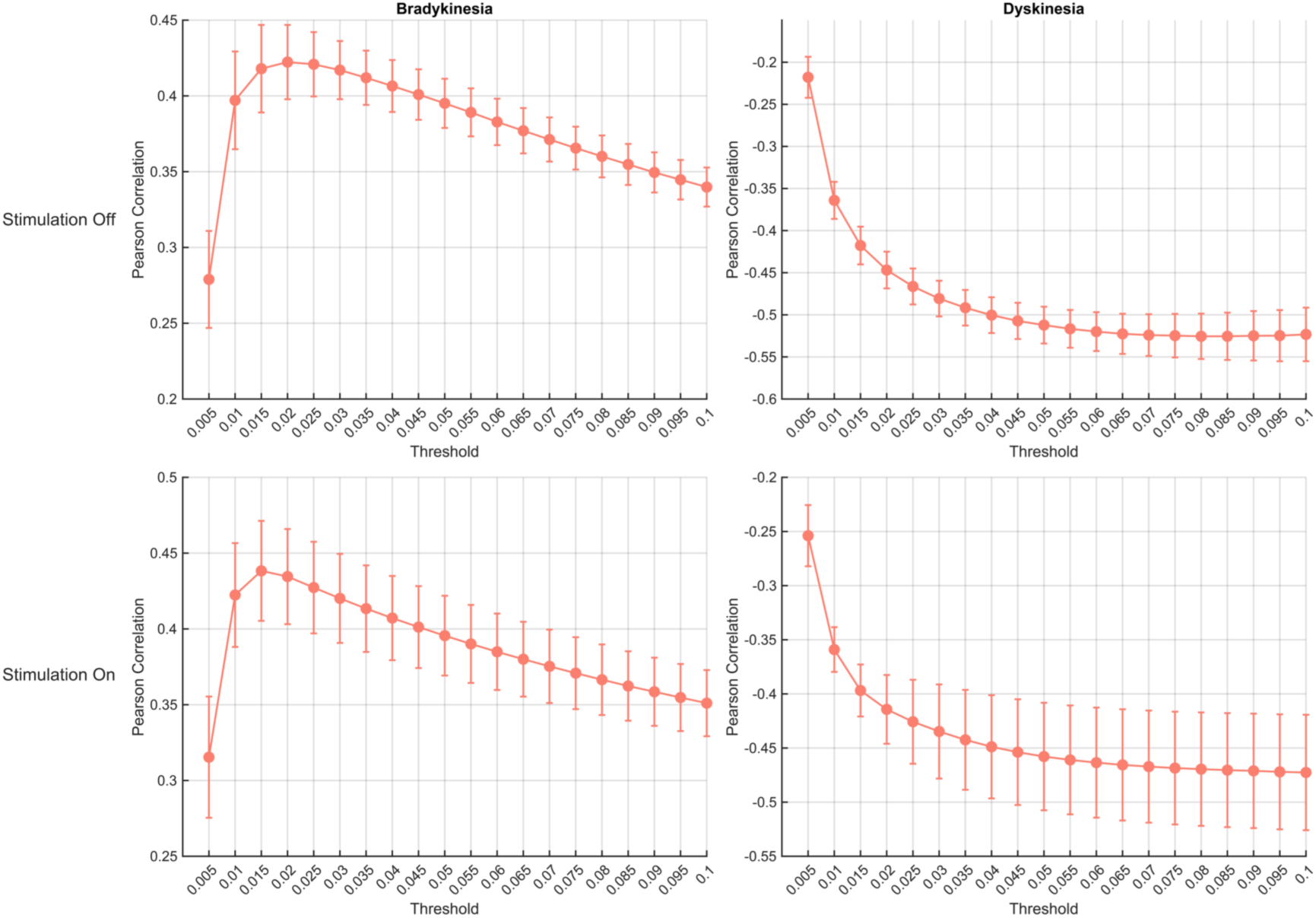
Grid search optimization for the time below threshold accelerometer feature. Threshold range, 0.005 to 0.1; step, 0.005. The Pearson correlation coefficient between time below threshold and wearable bradykinesia or dyskinesia scores is used for optimization. The threshold with the highest absolute coefficient was selected as the optimal threshold. For bradykinesia, optimal thresholds were 0.02 (stimulation off) and 0.015 (stimulation on). Similarly for dyskinesia, optimal thresholds were 0.08 (stimulation off) and 0.1 (stimulation on).

**Supplementary Figure 2:**
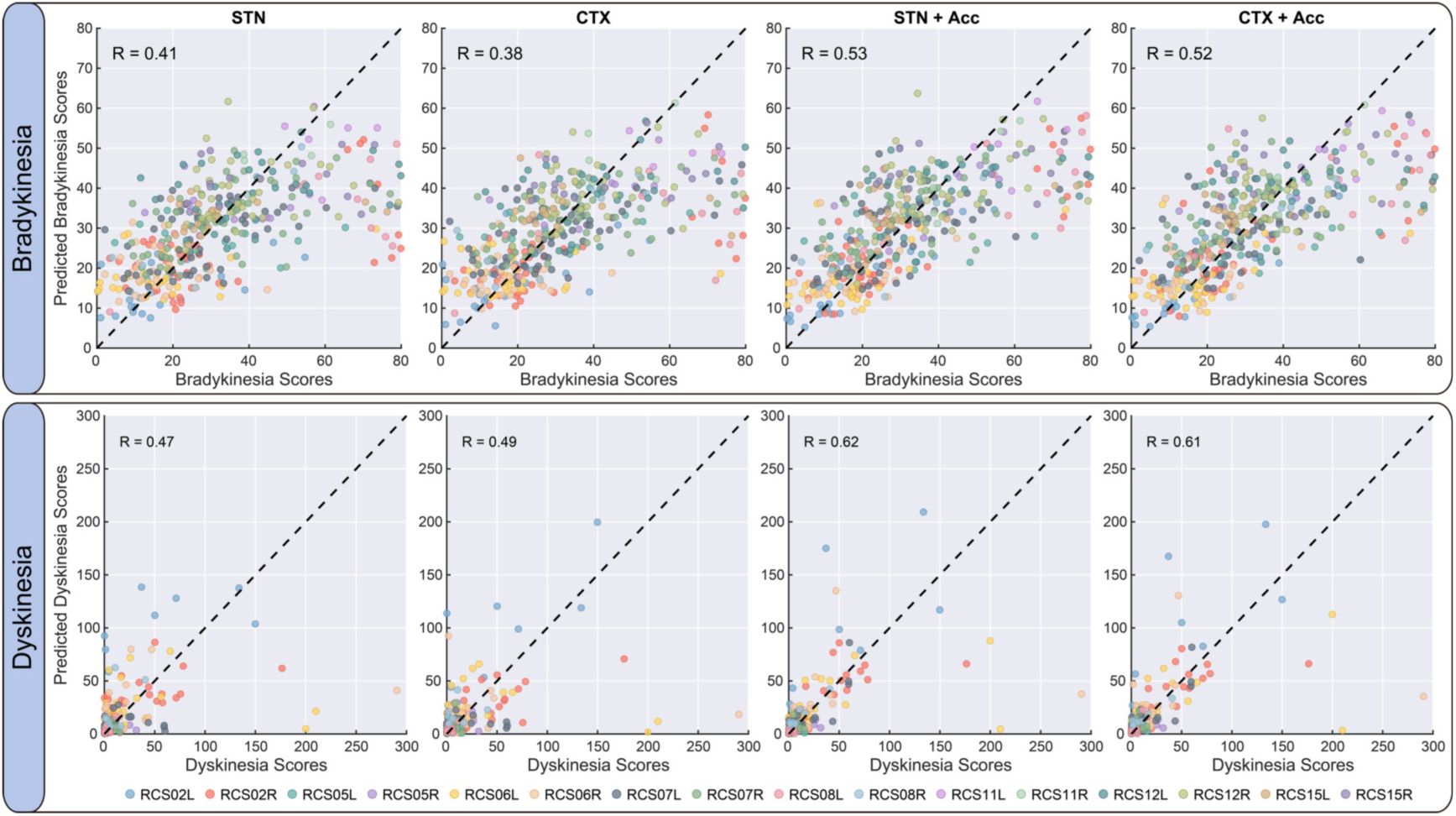
Performance of the random forest regressor at predicting PKG derived bradykinesia and dyskinesia severity from device accelerometer and neural activity recordings. Predicted vs. true bradykinesia and dyskinesia scores for each hemisphere (different colours indicate different hemispheres), shown separately for predictions using STN neural features, cortical neural features and the combination of each of these features with device accelerometer features. For visualisation purposes the plotted dots are downsampled from the validation set of each fold of five-fold cross-validation, in a ratio of 1:50. The coefficient of determination was computed for all data points. Points on the diagonal (black dotted line) indicate perfect predictions.

**Supplementary Figure 3:**
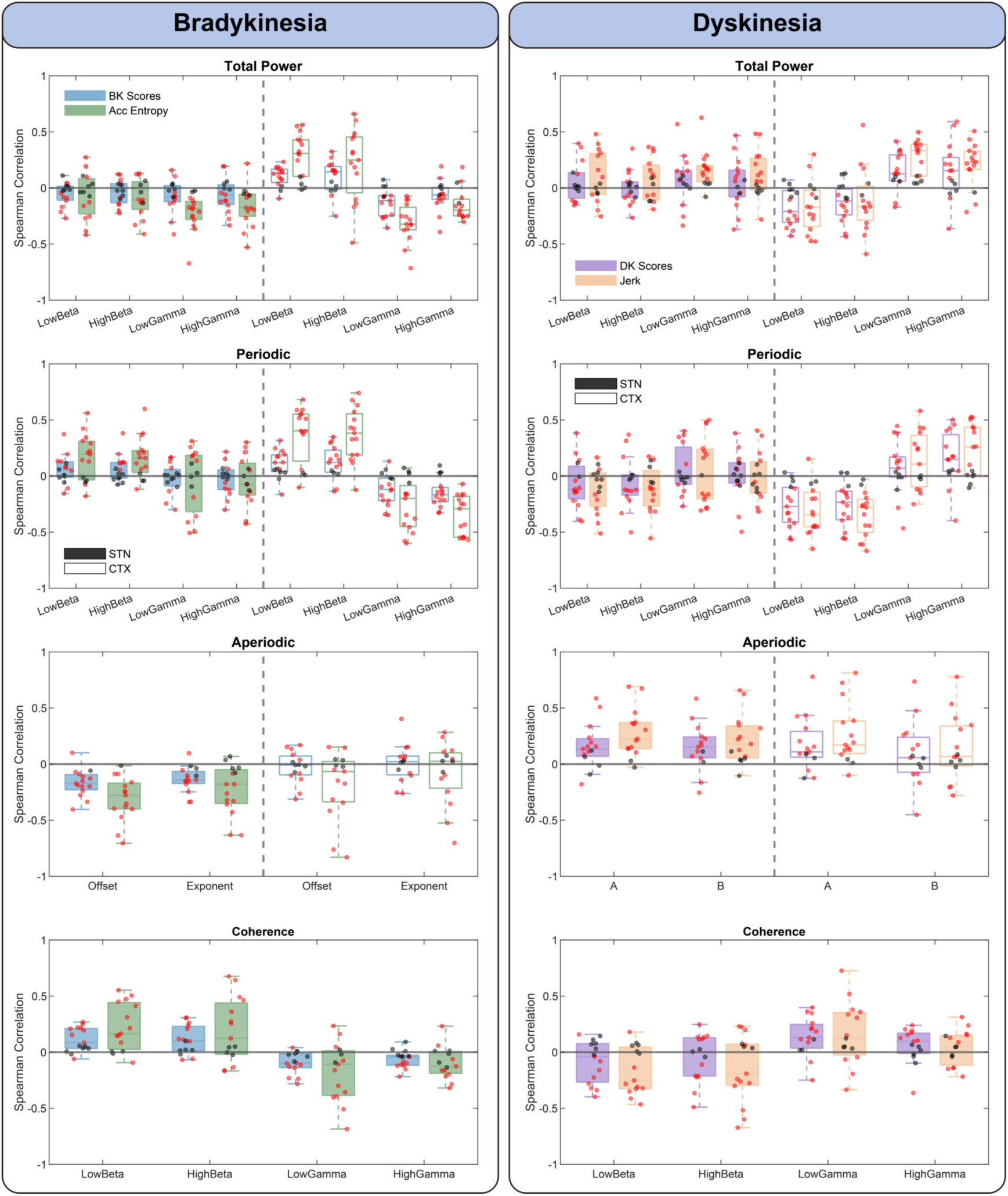
Optimal accelerometer features recapitulate neural correlates of motor symptom severity during continuous STN DBS. Hemisphere-wise Spearman correlations between neural features and either PKG-derived motor scores or the corresponding optimal accelerometry-derived surrogate. Colours within each subplot denote correlation type (PKG score vs. device accelerometry; see legend). Dark shaded panels indicate STN features, whereas light shaded panels indicate motor cortical (CTX) features. Red points indicate hemispheres with statistically significant correlations after Benjamini–Hochberg false discovery rate correction (q < 0.05), whereas black points indicate non-significant associations. Offset, aperiodic offset; exponent, aperiodic exponent. STN, subthalamic nucleus; CTX, motor cortex.

**Supplementary Figure 4:**
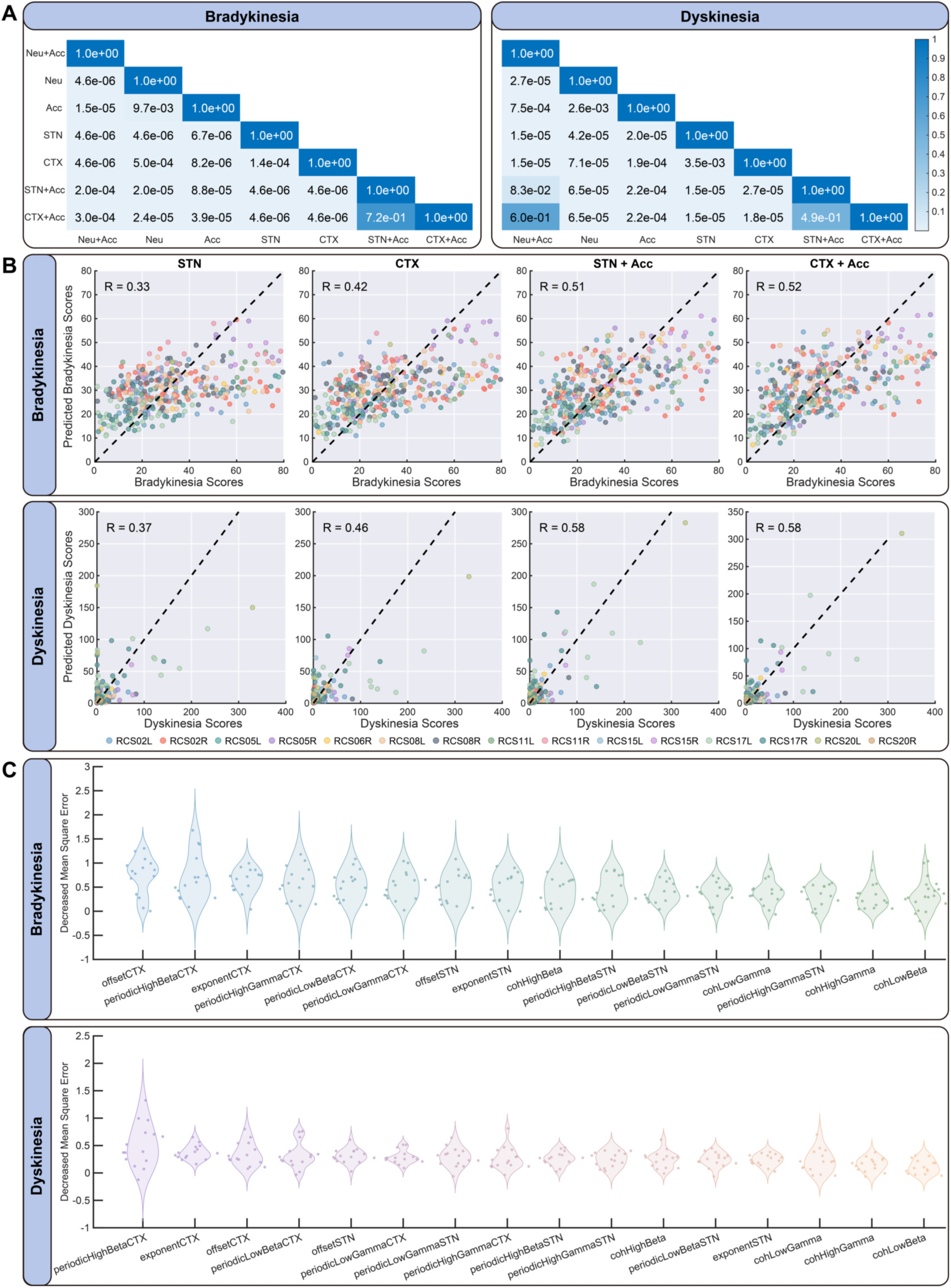
Device accelerometry outperforms neural activity features for predicting motor symptom severity during continuous STN DBS. Data are shown for the Random Forest regressor. **A:** Pairwise comparisons of feature sets for bradykinesia and dyskinesia prediction. P-values were computed using Wilcoxon signed-rank tests on mean R² values across cross-validation folds. Multiple comparisons were corrected using the Benjamini–Hochberg false discovery rate, and q-values are reported. Colour indicates q-value. **B:** Predicted versus observed bradykinesia and dyskinesia scores across hemispheres for models trained using STN neural features, cortical neural features and the combination of each of these features with device accelerometer features. For visualisation, a 1-in-50 random subsample of validation data is shown; R² values were computed using all observations. **C:** Feature importance for bradykinesia and dyskinesia prediction. Neural features are ordered by mean importance across hemispheres, quantified using out-of-bag (OOB) permutation importance. Each point represents one hemisphere.

